# PmaI-dependent pH gradients initiate inter- and intraspecies germling fusion in filamentous fungi

**DOI:** 10.1101/2025.06.24.661326

**Authors:** A. Y. Rudolph, N. Rotermund, N. Vasilevska, A. Fleißner, D.E. Nordzieke

**Affiliations:** University of Göttingen, GZMB, Institute of Microbiology and Genetics, Genetics of Eukaryotic Microorganisms, Grisebachstr. 8, 37077 Göttingen, Germany; University of Braunschweig, Institute of Technology, Institute of Genetics, Spielmannstraße 7, 38106 Braunschweig, Germany

**Keywords:** *Colletotrichum graminicola*, *Neurospora crassa*, *Botrytis cinerea*, PmaI ATPase, fungal GPCR, MAPK pathway, CAT fusion

## Abstract

Somatic cell fusion among spore germlings is a complex process in filamentous fungi that enables the formation of a syncytial unit and facilitates the exchange of genetic material apart from sexual reproduction. In this study, we investigated the molecular mechanisms underlying germling fusion in the maize pathogen *Colletotrichum graminicola*. Based on secretomes harvested during the germling fusion process of fusion-competent oval and fusion-deficient falcate conidia, we identified pH as a communication signal. We showed that the plasma membrane localized ATPase PmaI is a probable source for pH gradient formation. Our results also suggest that the a-pheromone receptor CgSte3 mediates sensing of acidic pH gradients, activating downstream the Cell Wall Integrity Mitogen Activated Protein Kinase pathway. We propose a new model for germling interactions, in which pH serves as an interaction initiation signal, resulting in the activation of the cell-dialogue. Our findings have implications for our understanding of fungal communication and pathogenicity, and highlight the importance of pH as a signal molecule in germling fusion.

## Introduction

Somatic cell fusion is a complex process that enables filamentous fungi to merge genetically identical or non-identical cells, resulting in the formation of a syncytial unit. This process is essential for the efficient development and growth of fungal colonies, and is mediated by a variety of signaling pathways and molecular mechanisms (Bastiaans *et al.*, 2015, Richard *et al.*, 2012, Herzog *et al.*, 2015). In *Neurospora crassa*, a model organism for studying fungal biology, somatic cell fusion is initiated by the germination of conidia, which undergo mutual attraction and fusion to form a syncytial unit (Read *et al.*, 2012). This process is regulated by a complex interplay of signaling molecules and molecular components, including transcription factors, the striatin-interacting phosphatase and kinase (STRIPAK) complex, NADPH oxidase complexes, and mitogen-activated protein kinase (MAPK) pathways (Fischer and Glass, 2019).

The current understanding of germling fusion in fungi suggests that it is a three-step process:

(I) the formation of conidial anastomosis tubes (CATs), (II) bidirectional chemotropic growth of CATs towards each other (CAT homing), and (III) cell fusion at the CAT contact side (Glass *et al.*, 2004, Kurian *et al.*, 2018). Central to the reciprocal recognition of interaction partners are the two proteins SO (soft) and MAK-2, which are part of the cell wall integrity (CWI) and pheromone Mitogen Activated Protein Kinase (MAPK) pathways, respectively (Fleissner *et al.*, 2009, Serrano *et al.*, 2018). During the tropic growth of the interacting cells, both proteins are recruited to the plasma membrane of the growing cell tips in an alternating manner. The current hypothesis proposes that SO localization to the CAT tip is associated with the release of a previously unknown communication signal, which is recognized by MAK-2 in the interaction partner, forming the so-called ‘cell-dialogue’ (Fleissner et al., 2009). Recent studies have challenged the long-held assumption that germling fusion occurs only among germlings of the same species, revealing that interspecies interactions and true fusion can occur between different fungal species (Haj Hammadeh *et al.*, 2022, Mehta and Baghela, 2021). For example, germlings of *N. crassa* and *Botrytis cinerea* interact with each other albeit the fusion process is not completed (Haj Hammadeh et al., 2022). Additionally, *Colletotrichum gloeosporioides* and *Colletotrichum siamense* fuse and form viable progeny, with the genome of the resulting progeny undergoing genetic recombination (Mehta and Baghela, 2021).

*Colletotrichum graminicola* is a hemibiotrophic maize pathogen responsible for anthracnose disease, causing high economic damage with corn loss of 10-20% of harvest annually (Belisário *et al.*, 2022). For successful disease development, *C. graminicola* generates two different asexual spore types, falcate conidia in acervuli on infected maize leaves, and oval conidia inside parenchyma cells nearby the plant vascular system (Belisário et al., 2022, Panaccione *et al.*, 1989). Those spore types differ significantly in their vegetative development and infection-relevant processes, making falcate conidia more efficient in leaf infection and enable oval conidia to sense maize root-exudated signals, followed by chemotropic growth towards the host and root infection (Nordzieke *et al.*, 2019, Schunke *et al.*, 2020, Rudolph *et al.*, 2024a, Chaky *et al.*, 2001). One prominent difference between both spore types is the ability to form a network by vegetative fusions: oval conidia form fusion bridges at high rates under starvation conditions, whereas falcate conidia stay dormant in the same experimental setups. However, close proximity of oval and falcate conidia can release dormancy in falcate conidia and result in fusion of both spore types (Nordzieke et al., 2019). From these results we conclude that the germling interaction signal is secreted by oval conidia but is absent from falcate conidia. Intriguingly, oval conidia fusion takes place also on leaves. There, fusion bridge formation is correlated to the generation of hyphopodia for leaf penetration, although both processes do not directly depend on each other (Nordzieke et al., 2019, Nordzieke, 2022).

In this study, we aimed to identify signals responsible for reciprocal germling attraction during the fusion process in *C. graminicola*. To identify these signal(s), a methodical setup was established and verified, enabling us to collect secreted molecules during the fusion process. Comparing the germling fusion secretome of oval and falcate conidia, we observed difference in secretome pH. In the following, we verified that distinct pHs are able to attract *C. graminicola* germlings and medium pH influences CAT fusion frequency. We further explored that the a-pheromone receptor CgSte3 and the CWI pathway mediate pH sensing. Applying an inhibitor of the plasma membrane standing ATPase PmaI, we provided evidence that pH signal formation is dependent on ATPase activity and links germination and fusion in *C. graminicola*. From those findings, we propose a new model for germling interactions, in which pH serves as an interaction initiation signal, resulting in the activation of the cell-dialogue. Since PmaI-dependent pH gradients are formed in all fungi in the course of germination, we further explored intraspecies interaction of *C. graminicola* with *N. crassa* and *B. cinerea*. All together, we identified that pH gradients serve as signal during reciprocal germling attraction prior to fusion in *C. graminicola*. Since the fusion rate of other filamentous fungi does depend on pH as well, we propose a general function as interaction initiation signal.

## Material and methods

### Strains, growth conditions, and collection of spores

The *C. graminicola* (Ces.) G.W. Wilson (teleomorph *Glomerella graminicola* D. J. Politis) wild-type strain M2 (also referred to as M1.001) was provided by H. Deising (Martin-Luther University Halle-Wittenberg) and used in this study (Forgey *et al.*, 1978, O’Connell *et al.*, 2012). Falcate and oval conidia of *C. graminicola* were generated and collected as described previously (Rudolph *et al.*, 2024b). *N. crassa* matA (strain FGSC2489 provided by Fungal Genetics Stock Center) and *B. cinerea* (strain B05.10; Quidde *et al.*, 1999) were grown on Vogel’s minimal medium (Vogel’s MM), supplemented with 2% sucrose (Vogel, 1956). All strains used in this study are listed in Supplementary Table 1.

### Generation and analysis of germling secretome during conidial interaction

In our standard protocol for the quantification of germling fusions, we spread 50 μl of spore solution (c = 5 × 10^7^ ml^−1^) on water agar (1% serva agar, 1% agarose, 25 mM NaNO_3_) and incubate the plates for 17 h at 23°C. For oval conidia preparations, we see a high percentage of germling interacting and fusing after that time. However, falcate conidia do not germinate nor fuse under these conditions (Nordzieke et al., 2019). To enable the identification of signals generated during the germling interaction and fusion process, we aimed to collect the secretome of oval and falcate in this experimental setup and to compare their chemical properties. Therefore, we spread oval and falcate conidia of *C. graminicola* CgM2 on cellophane (SERVA cellophane sheets II (140 × 133), SERVA Electrophoresis GmBH, Heidelberg, Germany; 50 μl of spore solution with c = 5 × 10^7^ ml^−1^) topping water agar and let the liquid evaporate (clean bench). Afterwards, the cellophane sheets were transferred to petri dishes filled with a single layer of glass beads (Glass Beads Ø 2.85 – 3.45 mm, Roth, Karlsruhe, Germany) and 10 ml of a 25 mM NaNO_3_ solution. After 17 h incubation at 23°C, the liquid was collected in glass ware and frozen at −80°C. To control the formation of germling fusion networks, microscopic pictures were taken, verifying the formation of fusion bridges for oval conidia and their absence for falcate conidia (Figure 1). pH of secretome solutions was measured using pH strips (20RO5121, Roth, Karlsruhe, Germany) of freshly harvested or 10 x concentrated secretome.

**Figure 1.**
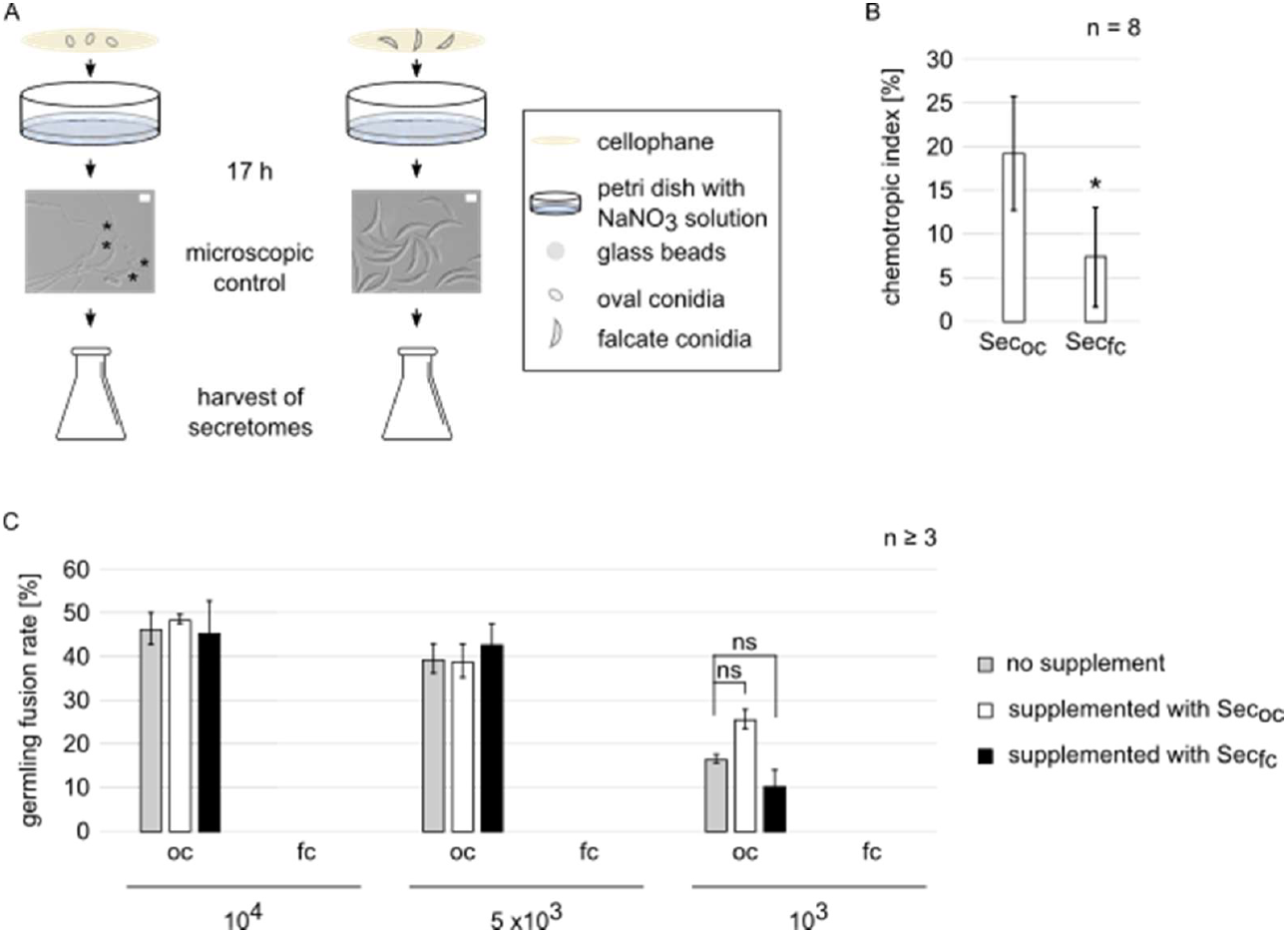
CAT secretome of *C. graminicola* oval conidia elicits chemotropic growth and shapes fusion and germination of oval and falcate conidia. (A) Schematic representation of fusion secretome generation. 50 μl of oval and falcate conidia spore solutions (c = 5 × 10^7^ ml^−1^) of the *C. graminicola* wildtype were spread on cellophane topping water agar. After evaporation of the liquid, cellophane sheets were transferred to petri dishes filled with a single layer of glass beads and 10 ml of a 25 mM NaNO_3_ solution. After 17 h incubation at 23°C, the fusion secretome was collected and the cellophane evaluated for the presence or absence of fusions. (B) Chemotropic index of *C. graminicola* germlings derived from oval conidia to 10 x concentrated fusion secretome harvested from oval (Sec_oc_) or falcate (Sec_fc_) conidia incubations. 20 μl of oval conidia (c = 5 × 10^5^ ml^−1^) were applied in a 3D printed device between two wells containing 40 μl solvent control and attractant. Microscopic evaluation of fungal tip growth was performed after incubation for 6 h, 23°C, n = 8 biological replicates, statistical significance was calculated with a two-tailed t-test (*, p < 0.05). (C) Germling fusion rates in dependence of spore concentration and the supplementation with 10 x concentrated Sec_oc_ and Sec_fc_. 10 μl of oval or falcate conidia were dropped in different amounts (10^4^, 5 × 10^3^, 10^3^) on water agar plates agar (1% serva agar, 1% agarose, 25 mM NaNO_3_). After drying, the drops were topped with 20 μl of 10 x concentrated Sec_oc_ and Sec_fc_ and the liquid dried again. After 17 h incubation at 23°C, at least 100 contacting germlings and conidia were microscopically examined for the formation of fusion bridges, n ≥ 3, ns = not significant.

### Germling fusion and germination assays in *C. graminicola*

To determine the impact of concentrated germling fusion secretome on the fusion and germination ability of oval and falcate conidia, 10 μl of different concentrated spores solutions (c = 10^6^ ml^−1^; 5 × 10^5^ ml^−1^; 10^5^ ml^−1^) were dropped on water agar (1% serva agar, 1% agarose, 25 mM NaNO_3_) and dried under a laminar flow. The dried spore drops were topped with 20 μl of 10 x concentrated secretome generated from oval or falcate conidia germlings during fusion conditions. After drying of the secretome drops, the plates were inoculated for 17 h at 23°C. At least 100 touching germlings and conidia were accessed for fusion bridge formation and 100 conidia for germination in at least three biological replicates.

For the determination of pH-dependent germling fusion and germination frequencies, 50 μl of oval conidia (c = 5 × 10^7^ ml^−1^) were spread on water agar (1% serva agar, 1% agarose, 25 mM NaNO_3_) adjusted to different pH values after autoclaving using pH strips (pH-Fix 4.5-10.0, Roth, Karlsruhe, Germany). For buffered conditions, the water agar was prepared with citrate phosphate buffer to reach a stable pH of 6.5. Microscopic evaluation was performed after 17 h of cultivation at 23°C, counting at least 100 touching conidia or germlings and accessing their fusion state. Germination rates were determined by evaluating at least 100 conidia. Both experiments were performed in at least three biological replicates.

Concentration-dependent germling fusion assays were performed by dropping 10 μl of oval conidia spore solution (10^6^ m^−1^; 7.5 × 10^5^ m^−1^; 5 × 10^5^ m^−1^; 2.5 × 10^5^ m^−1^; 10^5^ m^−1^; 7.5 × 10^4^ m^−1^; 5 × 10^4^ m^−1^) of *C. graminicola* wildtype (CgM2), *Cgste3* deletion and complementation strains on water agar (1% serva agar, 1% agarose, 25 mM NaNO_3_), dried under laminar flow, and incubated for 17 h at 23°C. For higher concentrations, at least 100 touching conidia or germlings were microscopically analyzed for the presence of fusions, for lower concentrations the maximal amount of conidia and germlings present in the 10 μl drop were evaluated. For each concentration, the experiments were performed at least three times.

To determine the effect of PmaI inhibitor Demethoxycurcumin (DMCM; Demethoxycurcumin ≥ 98% HPLC, Sigma-Aldrich, St. Louis, USA) on germling fusion and germination rate of oval conidia, DMCM was dissolved in DMSO. Working solutions were prepared just before use by further diluting with distilled water. As control, diluted DMSO concentration was used equalizing 15 μM of DMCM. 20 μl of different DMCM concentrations or the control were applied on dried 10 μl drops of oval conidia (c = 10^6^ ml^−1^) on water agar (1% serva agar, 1% agarose, 25 mM NaNO_3_) and dried as well under a laminar flow. Microscopic evaluation was performed after 17 h of cultivation at 23°C, counting at least 100 touching conidia or germlings and accessing their fusion state. Germination rates were determined by evaluating at least 100 conidia. Both experiments were performed in at least three biological replicates.

### Chemotropic assay

Chemotropic growth is defined as directed growth along a gradient of a given chemical stimulus, which can be positive (attractant signal) or negative (repellant signal). To display positive and negative chemotropic growth responses, the ‘chemotropic index’ is used as a standard for result depiction, indicating the difference of a preferred growth direction compared to a situation, in which neither attraction no repulsion occurs (Turrà *et al.*, 2015). In our lab, we developed a 3D printed device to easily monitor directed growth responses (Schunke et al., 2020). Using this device, we determined growth direction of fungal germlings derived from *C. graminicola* oval conidia of CgM2 (wildtype), Δ*Cgste3*, Δ*Cgso*, and the corresponding complementation strains after incubation for 6 h at 23°C. As attractant molecules, we evaluated 10 x concentrated secretome of oval and falcate conidia generated during the germling fusion process and potassium phosphate buffer adjusted to different pH values (pH 6, 7,, 7.5, 8). All experiments were performed in at least two biological replicates.

### Microscopy

Light (differential interference contrast (DIC)) was performed with the Axiolmager M1 microscope (Zeiss, Jena, Germany). Images were captured with the Photometrix coolSNAP HQ camera (Roper Scienific, Photometrics, Tucson, AZ, USA) and processed with the ZEISS ZEN software (version 2.3, Zeiss). Alternatively, images were taken with the Axio Observer microscope (Zeiss, Jena, Germany). Images were captured with sCMOS pco.panda 4.2 M (Excelitas PCO GmbH) and processed with the software 4-D MICROSCOPE FLMV 1.1 and 4-D MICROSCOPE Observer Z.1 (Schnabel et al., 1997).

### Inter- and intraspecies interaction studies

The interspecies interaction experiments were performed on Vogel’s MM (20 ml of 50x Vogel’s trace elements solution, 20 g Sucrose, 15 g Agar-Agar; Vogel, 1956). The interactions experiment with *C. graminicola* and *B. cinerea* was carried out at 23°C with an incubation time of 17 h. First, fungal spore solutions were adjusted to distinct concentrations (*B. cinerea*: c = 3.33 × 10^7^ ml^−1^; *C. graminicola*: c = 5 × 10^7^ ml^−1^). 300 μl of *B. cinerea* spore solutions were plated on Vogel’s MM medium and 50 μl of *C. graminicola* oval conidia added. Using a pipette tip, the spores from both species were mixed and spread onto the medium plate. For the interspecies interaction experiment between *C. graminicola* and *N. crassa*, 50 μl of a *C. graminicola* oval conidia spore solution (c = 5×10^7^ x ml^−1^) was first distributed on Vogel’s MM medium using glass beads and incubated for 14 h at room temperature. After that time, 300μl of *N. crassa* spore solution (c = 10^7^ ml^−1^) was added on those pre-incubated plates, dried and further incubated at 28°C for 3.5 - 4 h. The interspecies interaction rate was calculated based on several screen fields in which spores of both species were present. In total, ≥ 76 interactions were calculated in three biological replicates.

pH-dependent fusion and germination rates of *C. graminicola, B. cinerea*, and *N. crassa* was performed on Vogels’ MM adjusted to distinct pH without buffering the medium. For *C. graminicola*, 50 μl of oval conidia spore solution (c = 5×10^7^ x ml^−1^) was spread on Vogels’ MM using glass beads. For *B. cinerea* and *N. crassa*, 300 μl of c = 10^7^ ml^−1^ spore solutions of were added on Vogels’ MM and spread using pipette tips. After incubation (*C. graminicola*: 17 h, 23°C, *B. cinerea*: 17 h, 23°C, *N. crassa*: 3 h, 30°C), at least 100 contacting germlings or conidia were accessed for formed fusion bridges. In the same experiments, germination rates were determined from at least 100 conidia. Both fusion and germination experiments were conducted in at least 3 biological replicates.

Localization of MAK-2-GFP was monitored during *N. crassa* – *N. crassa* and *N. crassa* – *C. graminicola* interactions, using the same conditions described above. As readout served fluorescence microscopy.

### Statistics

For statistical analysis in this study, the T-test for unequal variances was used for all experiments (Ruxton, 2006).

### AI statement

The AI chatbot ‘Chat AI’ provided by the GWDG at the Georg-August University Göttingen was used to improve the text of this manuscript regarding English grammar and typing.

## Results

### Molecules secreted from oval conidia during fusion induce chemotropic growth of *C. graminicola* germlings

While numerous proteins and second messengers are necessary for CAT fusion (Fu *et al.*, 2011, Fischer and Glass, 2019), signals resulting in bidirectional attraction of germlings are unknown. To explore the potential of CAT signal identification, we established an experimental design enabling to harvest secreted signals during fusion bridge formation. We spread equal amounts of fusion competent oval or fusion-incompetent falcate conidia on cellophane sheets topping water agar. After drying the spore solution, we transferred the cellophane to a new petri dish filled with a single layer of glass beads and 25 mM NaNO_3_ solution. After 17 h of incubation, the germination and fusion state of the cultures were evaluated, showing that oval conidia germinated in high percentages and undergo fusions eagerly. In contrast, falcate conidia did not germinate or form fusion bridges (Figure 1, A). Since those results are in line with previous observations (Nordzieke et al., 2019), glass beads and liquid were separated by filtration and the flow through analyzed.

In a first experiment, we tested whether the fusion secretomes of oval and falcate conidia are able to attract *C. graminicola* germlings using a 3D printed device developed in our group (Schunke et al., 2020). Indeed, we found that 10x concentrated fusion secretome from oval conidia (Sec_oc_) induce a strong chemotropic response, whereas the 10x concentrated fusion secretome of falcate conidia (Sec_fc_) showed minor attraction potential (Figure 1, B), indicating a successful harvest of interaction signal(s). In a second approach, we supplemented oval and falcate conidia with 10x Sec_oc_ and Sec_fc_ and accessed their germination and fusion rates after 17 h of incubation. Since germling fusion is a spore concentration-dependent process, we tested the effect of the secretomes on optimal (10^4^) and suboptimal (10^3^) spore concentrations (Figure 1, C, Supplementary Figure S1). For oval conidia, we observed a spore-density dependent effect of the secretomes: whereas in optimal experimental conditions, no effect of secretome addition was visible, 10x Sec_oc_ did increase the fusion frequency of oval conidia applied in suboptimal concentrations (Figure 1, C). Regarding falcate conidia, the addition of concentrated Sec_oc_ or Sec_fc_ did not induce fusions; however, in experimental conditions with 10^3^ applied spores, germination of this spore type was induced (Supplementary Figure S1). Together, these first experiments indicate that isolation of a CAT fusion signal was successful with the experimental approach chosen and point to a possible interconnection between germination and fusion.

### Distinct pHs attract *C. graminicola* germlings and induce germling fusion

Next, we aimed on secretome characterization. When we monitored the pH of the Sec_oc_ and Sec_fc_ and compared it with control conditions, we observed a more basic pH for 10x Sec_oc_ (Figure 2, A). We thus aimed to explore, whether pH can act as attractant for *C. graminicola* and influence fusion frequency. As depicted in Figure 2, B, pH of 7.5 strongly attracts germlings, whereas higher or lower pH values do not, indicating the sensing is specific to distinct pH gradients. Monitoring of CAT fusion events on water agar plates with adjusted pH values revealed that in comparison to not adapted conditions, pH values of 5 and >7 are inhibiting fusion bridge formation. In contrast, medium adjusted to pH 6.5 induces fusion frequency. From these results, we were wondering whether pH could be a mere factor of wellbeing for *C. graminicola* germlings, therefore enabling increased number of fusion events. To test this, we generated buffered water agar medium with a pH adjusted to 6.5, which provides optimal fusion condition in not buffered conditions. Intriguingly, no fusions were detected in those experiments, underpinning the necessity of active pH change by fungal germlings or conidia for fusion bridge formation (Figure 2, C). In the same experiments, we also monitored germination. In contrast to network formation by fusion, the germination process is only affected under buffered conditions (Supplementary Figure 2).

**Figure 2.**
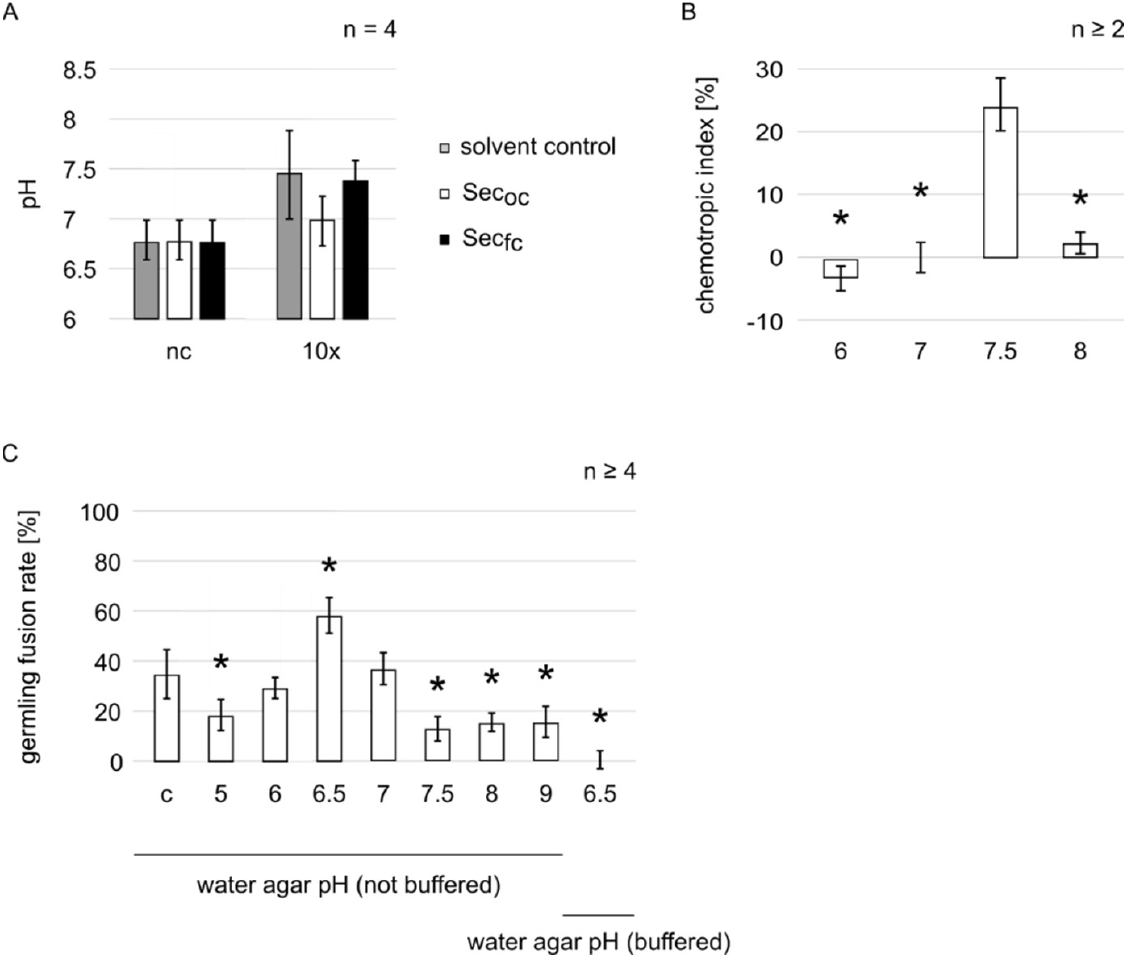
Distinct pH values attract *C. graminicola* germlings and impact their ability to undergo fusions. (A) pH of freshly harvested not-concentrated and concentrated fusion secretome of oval (Sec_oc_) and falcate (Sec_fc_) was examined using pH strips, n = 4. (B) Chemotropic index of *C. graminicola* wild-type germlings to potassium phosphate buffer adjusted to different pH values (pH 6; 7;7.5; 8). 20 μl of oval conidia (c = 5 × 10^5^ ml^−1^) were applied in a 3D printed device between two wells containing 40 μl solvent control and attractant. Microscopic evaluation of fungal tip growth was performed after incubation for 6 h, 23°C for least two biological replicates, statistical significance was calculated with a two-tailed t-test (*, p < 0.05). (C) Germling fusion rate of *C. graminicola* wild-type germlings spread on water agar (1% serva agar, 1% agarose, 25 mM NaNO_3_) adjusted to different medium pH with and without buffering. 50 μl of oval conidia (c = 5 × 10^7^ ml^−1^) were spread on the corresponding medium plates. After 17 h of incubation at 23°C, at least 100 contacting germlings and conidia were microscopically examined for fusion bridge formation, n ≥ 4. Statistical significance was calculated with a two-tailed t-test corresponding to control conditions (c) *, p < 0.05.

### The a-pheromone receptor CgSte3 and the MAPK scaffold CgSo mediate pH sensing

In recent work of our group and others, fungal pheromone receptors were shown to perceive defense molecules secreted from plant roots like peroxidases and diterpenoids besides fungal pheromones (Rudolph et al., 2024a, Rudolph et al., 2024b, Turrà et al., 2015, Sridhar *et al.*, 2020, Sharma *et al.*, 2022, Ramaswe *et al.*, 2024, Vitale *et al.*, 2019). Similarly, the Cell Wall Integrity Mitogen Activated Protein Kinase pathway (CWI) was shown to mediate the extracellular signals and transform them into chemotropic growth responses (Turrà et al., 2015, Sridhar et al., 2020, Sharma et al., 2022). To test whether this receptor-MAPK module is also involved in pH sensing, deletion mutants of genes encoding for the a-pheromone receptor *Cgste3* and the CWI scaffold *Cgso* were examined for their ability to sense and re-direct growth to pH 7.5 (Figure 3). In contrast to wild-type strain CgM2, which show a strong growth re-direction, both mutant strains fail to grow towards a gradient of pH 7.5. For both mutants, this effect was restored by reintroducing the corresponding gene (Figure 3, A).

**Figure 3.**
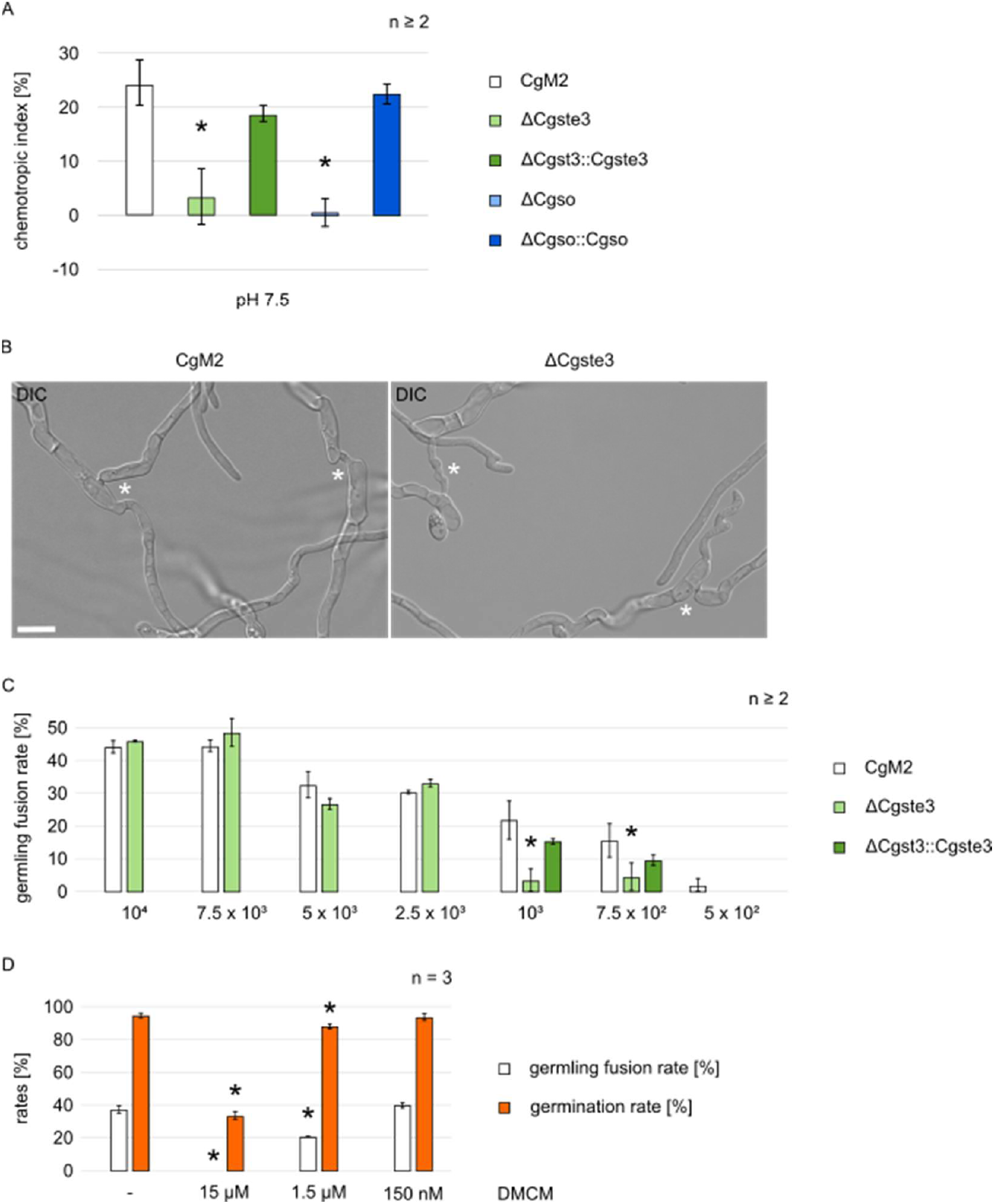
Sensing of acidic pH is mediated by the a-pheromone receptor CgSte3 and the CWI MAPK scaffold protein CgSo. (A) Chemotropic index of *C. graminicola* germlings of wildtype (CgM2), *Cgste3* and *Cgso* deletion strains and corresponding complementations to potassium phosphate buffer adjusted to pH 7.5. 20 μl of oval conidia (c = 5 × 10^5^ ml^−1^) were applied in a 3D printed device between two wells containing 40 μl solvent control and attractant. Microscopic evaluation of fungal tip growth was performed after incubation for 6 h, 23°C for least two biological replicates, statistical significance was calculated with a two-tailed t-test to CgM2 (*, p < 0.05). (B) Microscopic evaluation of germling fusions after spreading of 50 μl *C. graminicola* wildtype (CgM2) or Δ*Cgste3* oval conidia spore solutions (c = 5 × 10^7^ ml^−1^) on water agar (1% serva agar, 1% agarose, 25 mM NaNO_3_) and incubation for 17 h, 23°C, scale = 10 μm, fusions are indicated with white asterisk. (C) Spore concentration-dependent germling fusion rates of *C. graminicola* wildtype, Δ*Cgste3* and Δ*Cgste3* complementation strain Δ*Cgste3*::*Cgste3*. 10 μl of oval conidia spore solutions were dropped on water agar (1% serva agar, 1% agarose, 25 mM NaNO_3_) to reach spore amounts of 10^4^ - 5 × 10^2^. Microscopic evaluation after 17 h of incubation at 23°C was performed for at least 100 contacting germlings and conidia in higher spore concentration. In lower spore concentrations, the full area was examined for fusion bridge formation, n ≥ 2, statistical significance was calculated with a two-tailed t-test to CgM2 values of the corresponding spore concentration (*, p < 0.05). (D) Germling fusion and germination rates of *C. graminicola* wildtype in dependence of treatment with the PmaI inhibitor Demethoxycurcumin (DMCM). 20 μl of different DMCM concentrations or the control were applied on dried 10 μl drops of oval conidia (c = 10^6^ ml^−1^) on water agar (1% serva agar, 1% agarose, 25 mM NaNO_3_) and dried prior to incubation for 17 h at 23°C. Since DMCM was dissolved in DMSO, diluted DMSO equalizing 15 μM DMCM solutions were used as control. Germling fusion was accessed for at least 100 contacting conidia or germlings. Germination rates were determined by evaluating at least 100 conidian, n = 3, statistical significance was calculated with a two-tailed t-test to control conditions (-), *, p < 0.05.

In an earlier report, we investigated the impact of *Cgso* deletion on developmental processes in *C. graminicola*. Besides the absence of germling and hyphal fusion, this mutant shows a strong conidiation phenotype, provoked by the impaired nutrition of asexual fruiting bodies, the acervuli (Nordzieke, 2022). Therefore, Δ*Cgso* appear dark and generate increased levels of aerial mycelia compared to wildtype. In contrast, the a-pheromone receptor mutant Δ*Cgste3* shows normal level of conidiation, indicating that this strain is able to undergo vegetative fusions. Indeed, microscopic investigations showed normal fusion frequencies of Δ*Cgste3* compared to wildtype, when both strains are plated in optimal spore concentrations (Figure 3, B and C). Interestingly, a decrease in spore concentration revealed a significant reduction of Δ*Cgste3* fusion rate compared to wild-type CgM2, which was reversed in a *Cgste3* complementing strain. At the same time, Δ*Cgste3* germination rates don’t differ from the wildtype (Figure S3). From these results, we concluded that pH might act as long-distance signaling molecule, able to guided conidia or germlings towards each other, but dispensable when germlings are in close proximitiy.

Next, we aimed to identify the source of pH gradient formation. In filamentous fungi and plants, growth is strictly correlated to the activity of a plasma membrane located ATPase, PmaI. This

ATPase pumps protons into the extracellular space, creating pH gradients used for the uptake of diverse molecules (Portillo, 2000, Palmgren, 2001, Li *et al.*, 2022, Zhgun *et al.*, 2020). Since *pmaI* is an essential gene in many organisms (Harris *et al.*, 1994), we decided to use a Demethoxycurcumin (DMCM), a described inhibitor of PmaI (Dao *et al.*, 2016), to test a probable impact of PmaI-dependent pH gradients for germling fusion. Intriguingly, we see a concentration-dependent inhibition of germling fusion compared to control conditions (Figure 3 D). At the same time, germination is inhibited by DMCM, further indicating that germination and fusion are interconnected processes.

### Indications for pH as a conserved fungal fusion initiation signal

At first, germling fusion was considered as a highly species-specific process, providing only germling interaction and fusion of individuals of the same species or communication group (Heller *et al.*, 2016, Serrano et al., 2018). However, this perspective is changing. For filamentous fungi like *N. crassa* and *B. cinerea*, intraspecies germling interaction and even the induction of the SO-MAK-2-dependent cell dialog was shown (Haj Hammadeh et al., 2022). However, those interactions never result in cross-species progeny. In *N. crassa* – *B. cinerea* interactions, the fusion process is abolished upon contact of both interaction partners. Instead of cell wall degradation to allow fusion, a cell wall thickens (Haj Hammadeh et al., 2022). Additionally, there are reports about true fusions in between closely related species. The plant pathogenic fungi *Colletotrichum gloeosporioides* and *Colletotrichum siamense* form progeny after fusion, which show signs of genetic recombination of both parental species, making intraspecies fusion a driver of evolution and host adaptation (Mehta and Baghela, 2021, Haj Hammadeh et al., 2022). A prerequisite for intraspecies fusion is the presence of common or very similar interaction signals, which could guide individuals of two different species towards each other. Since PmaI-dependent pH gradients are formed by all fungi during conidial germination, we were interested in testing, whether *C. graminicola* germlings are able to interact with other fungal species.

In a first approach, we plated either fusion competent spores of *C. graminicola* and *B. cinerea* or *C. graminicola* and *N. crassa* on Vogel’s minimal medium, providing conditions for fusion bridge formation in all three species. Since those fungi grow with different speed and have distinct temperature optima, specific incubation times had to be chosen. For *C. graminicola* – *B. cinerea* interactions, conidia were spread in different concentrations on Vogel’s MM and incubated at 23°C. Since *N. crassa* conidia germinate and fuse much faster than *C. graminicola*, the latter was pre-incubated for 14 h. After that time-interval, *N. crassa* conidia were added and both fungi incubated further at 28°C. Both described experimental setups resulted in interspecies and intraspecies interactions, whereas the proportion of species-species interactions was least (Figure 4, A - D).

**Figure 4.**
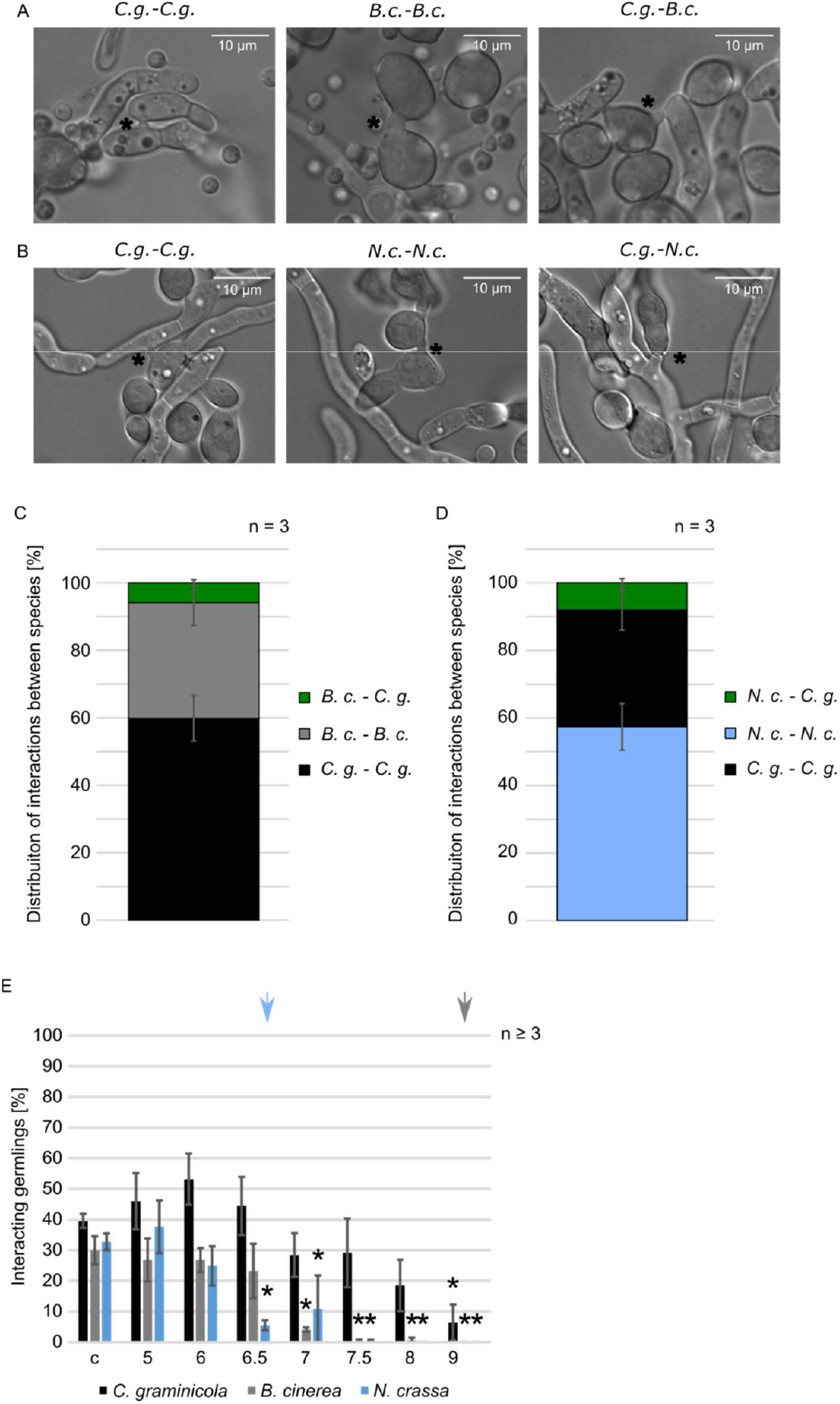
Interaction of *C. graminicola* with *N. crassa* and *B. cinerea*. (A-D) Inter- and Intraspecies interactions of *C. graminicola* and *B. cinerea* or *N. crassa* on Vogels minimal medium (MM). (A, C) The interactions experiment with *C. graminicola* and *B. cinerea* was carried out at 23°C with an incubation time of 17 h. First, *C. graminiocla* oval conidia (50 μl of c = 5 × 10^7^ ml^−1^) and 300 μl *B. cinerea* spore solution (c= 3.33 × 10^7^ ml^−1^) were spread together on Vogels’ MM. (B, D) To access *C. graminicola* and *N. crassa* germling interactions, 50 μl of a *C. graminicola* oval conidia spore solution (c = 5×10^7^ x ml^−1^) was distributed on incubation medium and incubated for 14 h at 23°C. After that time, 300μl of *N. crassa* spore solution (c = 10^7^ ml^−1^) was added on those pre-incubated plates and further incubated at 28°C for 3.5 - 4 h. (A-B) Microscopic images of intra- and interspecies interaction, interactions are marked with a black asterisk, scale = 10 μm. (C-D) Inter- and intraspecies interaction rates of ≥ 76 germling interactions of *C. graminicola* and *B. cinerea* (C) and *C. graminicola* with *N. crassa* (D), n = 3. (E) Germling interaction rates of *C. graminicola, B. cinerea* and *N. crassa* on Vogel’s minimal medium adjusted to different pH. Species-specific conidia amounts (*C. graminicola*: 50 μl of 5 × 10^7^ ml^−1^; *B. cinerea* and *N. crassa*: 300 μl of c = 10^7^ ml^−1^) were spread on Vogels’ MM and incubated at 23°C for 17 h (*C. graminicola, B. cinerea*) or 28°C for 3,5 - 4 h (*N. crassa*), n ≥ 3, statistical significance was calculated with a two-tailed t-test to control conditions (c), *, p < 0.05.

The hallmark of germling interaction in *N. crassa* is the oscillatory cell-dialogue, in which SO and MAK-2 show an alternate recruiting to the plasma membrane of interacting partners. At the end of this dialogue, MAK-2 accumulates at the contact point. Intriguingly, in intraspecies interactions with *N. crassa*, this late accumulation of MAK-2 was shown, indicating an intact cell dialogue (Fleissner et al., 2009, Haj Hammadeh et al., 2022). To test whether this is true for *N. crassa* – *C. graminicola* interactions as well, a *N. crassa* wildtype strain expressing MAK-2-GFP was used in interaction studies with *C. graminicola*. Likewise as described for *N. crassa* – *N. crassa* and *N. crassa* – *B. cinerea* interactions, MAK-2-GFP accumulates at the contact point with *C. graminicola* germlings (Supplementary Figure 4).

Together these results indicate, that indeed all three fungi share a common interaction signal, but indicate also species-specific signals, explaining the higher proportion of self-interactions observed. To test whether the shard interaction signal might be pH, conidia of all three species were grown on Vogel’s MM adjusted to different pH without buffering the medium. As seen for *C. graminicola* on water agar, interaction frequency of all three species is pH dependent, although to different extents: *C. graminicola* shows the most pH-stable interaction pattern, reduced only significantly at basic pHs (Figure 4 E). The interaction frequency of *B. cinerea* already dropped at pH 7, and in *N. crassa*, pH of 6.5 already reduced interaction rates significantly. A similar pattern can be seen for the germination of conidia on Vogel’s MM. Germination rates of *C. graminicola* oval conidia is hardly affected also at high pH and the germination rates of *B. cinerea* are stable until pH 9. However, *N. crassa* conidia are highly sensitive to pH changes, showing already significant reduction on medium pH of 6.5 (Supplementary Figure 5).

**Figure 5.**
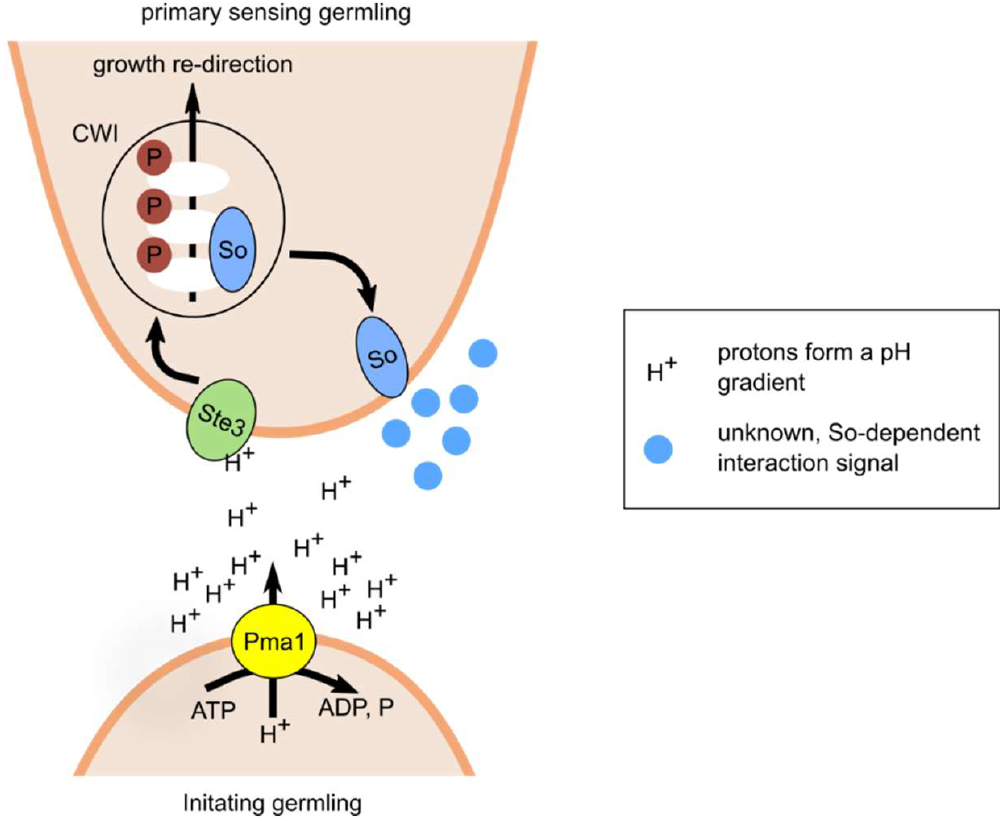
Working model for the pH-dependent induction of the cell dialogue. PmaI-depenent pH gradients (H^+^) form immediately when a conidium starts to germinate. Those pH gradients are sensed by surrounding germlings or conidia via the a-pheromone receptor Ste3, resulting in the activation of the Cell Wall Integrity (CWI) MAPK pathway and induction of growth redirection towards the interaction partner. From work in other fungi, we propose that CWI activation by Ste3 induces the relocalization of SO to the tip of the interacting germling, followed by the release of a SO-dependent, so far unknown communication signal (blue circles).

## Discussion

The perception of extracellular signals is important for fungi to find nutrients, hosts, toxic environments, or germlings and spores of the same species (Turrà *et al.*, 2016). In this study, we identified pH as a communication signal outgoing from secretomes harvested during germling fusion. Already in earlier studies, pH was identified as a fusion-modulation factor: over a broad pH range fusions *Fusarium oxysporum* take place, however, when the medium is buffered, only pH of 6.4 still allows fusion to occur (Kurian et al., 2018). From those results, it was concluded that an active modulation of pH by the fungal germlings are required for germling fusion. In our experiments, we see a similar influence on pH of fusion frequency of *C. graminicola*: whereas not-buffered media allow fusions over a broad pH spectrum, buffering abolishes fusion bridge formation. Analysis of chemotropic growth further emphasized the role of distinct pHs (7.5) on germling attraction and thus their signaling nature (Figure 2). Likewise, acidic pH was shown to induce chemotropic growth in germlings of *Aspergillus nidulans* (Yamamoto *et al.*, 2024). We further identified the plasma membrane localized ATPase PmaI as probable source for pH gradient formation using the specific PmaI-inhibitor Demthoxycurcumin (Dao et al., 2016). In fungi and plants, PmaI ATPase-formed pH gradient drive the uptake of nutrients and its activity is directly linked to cell growth (Serrano, 1984, Divon and Fluhr, 2007, Yamamoto et al., 2024, Palmgren, 2001, Haruta *et al.*, 2010, Morsomme and Boutry, 2000). Additionally to the provision of an extracellular proton gradient, PmaI activity results in a lower cytosolic pH, which is involved in the regulation of MAPK pathways and thus affects development and pathogenicity in *F. oxysporum* (Mariscal *et al.*, 2022). Intriguingly, PmaI was also shown to be crucial for the sensing of acidic pH in *A. nidulans*, however it is unclear how the determining mechanism works (Yamamoto et al., 2024).

pH is a crucial environmental factor for fungi and its fluctuations influence diverse cellular processes like cell wall remodeling, nutrient availability, and enzyme activity (Li *et al.*, 2021). Many plant and human pathogenic fungi modulate the extracellular pH to explore their surroundings, enable successful host infection and maintain cellular homeostasis (Masachis *et al.*, 2016, Fernandes *et al.*, 2017, Kane, 2016, Alkan *et al.*, 2013, Vylkova, 2017, Kesten *et al.*, 2019). Traditionally, plant-interacting fungi were thought to be strictly alkalizing or acidifying their surroundings. However, in the last decade it was shown, that induced changes in ambient pH are flexible and depend on carbon availability (Bi *et al.*, 2016). Sensing of extracellular, alkaline pH is primarily mediated by the pal/RIM pathway in fungi, with PalH as the central pH sensor (Selvig and Alspaugh, 2011). Together with MAPK modules, the pal/RIM pathway is central for the sensing and the response to extracellular pH levels, enabling to maintain cellular integrity also in the face of stress and pathogenic host interaction (Lara-Martínez *et al.*, 2025). PalH is a 7 transmembrane receptor resembling classical G-protein coupled receptors (GPCR), interacting downstream with an arrestin for signal transduction (Lucena-Agell *et al.*, 2016). As transcription factor downstream of PalH, PacC is described, acting as activator of gene expression at alkaline pH and as a repressor of genes expressed at acidic pH (Lara-Martínez et al., 2025). In contrast, the sensor for acidic environmental pH is unknown in fungi (Yamamoto et al., 2024). In this study, we provide evidence that the a-pheromone receptor CgSte3 mediates sensing of acidic pH gradients, activating downstream the CWI pathway (Figure 3). Fungal pheromone receptors are GPCRs first described in the yeast *Saccharomyces cerevisiae*, in which they mediate chemotropic attraction of mating partners during sexual fusion (Hagen and Sprague Jr, 1984, Nakayama *et al.*, 1985, Jenness *et al.*, 1986). In the last years, new roles for fungal pheromone receptors were identified. This includes the spore density dependent regulation of germination in *F. oxysporum*, and the sensing of plant defense molecules like class III peroxidases and diterpenoids in different host-pathogen systems (Turrà et al., 2015, Vitale et al., 2019, Sharma et al., 2022, Rudolph et al., 2024a, Rudolph et al., 2024b). Downstream of diterpenoid and peroxidase sensing, in those studies the CWI was shown to be responsible for consequent growth redirection (Turrà et al., 2015, Sharma et al., 2022, Rudolph et al., 2024b), indicating a conserved link between external signal sensing by pheromone receptors and CWI activation. So far, only one study investigated how the two pheromone receptors interact upon peroxidase sensing. These results point to a scenario, in which peroxidase directly or a peroxidase-generated ligand stimulates heterodimer formation in between Ste2 and Ste3 of *Fusarium graminearum* (Sharma et al., 2022). Intriguingly, *C. graminicola* is the only species, in which the α-pheromone receptor Ste2 was lost during evolution (Wilson *et al.*, 2021) and therefore the recognition of external signals is dependent only on the a-pheromone receptor Ste3. How the perception in this one-receptor system functions on the molecular level, is still to be explored. Another open question is how those different attractant molecules are sensed by the same receptor. Since the chemical properties of the perceived signals are such diverse, a direct ligand-receptor interaction seems unlikely. For mammalian cells it was shown that GPCRs and growth factor receptor tyrosine kinases (RTKs) can activate each other in a process termed ‘transactivation’, without the requirement for ligand binding to the secondary receptor (Wang, 2016, Delcourt *et al.*, 2007). In such a scenario, the a-pheromone receptor CgSte3 could be associated with other, more specific receptors, which serve as sensors for diterpenoids, peroxidases or pH specifically. Additionally, modifications of amino acids can induce GPCR activation (Wang *et al.*, 2018). In mammalian cells, a current study shows that protonation of histidine residues can induce GPCR activation (Guo *et al.*, 2025). As a third possibility, pH could influence the binding-capacity of another ligand by CgSte3, resulting in receptor activation (Sun *et al.*, 2025). Which of these possibilities or also other probable mechanisms are mediating CgSte3 receptor activation, has to be investigated in future studies.

Together, our results about *C. graminicola* CAT fusion point to a scenario, in which PmaI-depenent pH gradients are formed as soon as a conidium starts to germinate. Those pH gradients are sensed by surrounding germlings or conidia via the a-pheromone receptor CgSte3, resulting in the activation of CWI and induction of growth redirection towards the interaction partner. From work in other fungi, especially *N. crassa*, we know the oscillatory recruitment of SO and MAK-2 during germling interaction is crucial for fusion bridge formation (Fleissner et al., 2009, Serrano et al., 2018). Since CgSo is a scaffold of CWI, pH sensing might additionally induce CgSo localization to the plasma membrane, which could be associated with the secretion of a second interaction signal, still to be discovered (Figure 5). This model could also explain why germling fusions are absent in falcate conidia: those spores have a pronounced dormant phase, mediated by secondary metabolites like mycosporines and other factors (Leite and Nicholson, 1992, Vasselli *et al.*, 2022, Chaky et al., 2001). Since germination of falcate conidia does not occur under starvation conditions, PmaI is inactive and no pH gradients build up, making an induction of the fusion process impossible. However, when oval conidia are close by, they can provide the pH fusion-initiation signal, which is perceived by falcate conidia and subsequently induce dormancy breaking, germling interaction, and fusion.

The ability of germlings to fuse in an early stage of colony formation, fastens radial growth, by sharing of nutrients or cellular components (Roca *et al.*, 2005, Richard et al., 2012). Recently, studies have shown that germling communication mediating bidirectional chemotropic growth prior to fusion is not species-specific (Haj Hammadeh et al., 2022). Closely related species of *Colletotrichum* and *Aspergillus* are even able to fuse and exchange genetic material, resulting in progeny with recombinant genomes (Ishitani *et al.*, 1956, Mehta and Baghela, 2021). In this study, we show that the maize pathogen *C. graminicola* communicates with the saprophyte *N. crassa* and the pathogen *B. cinerea* with 8 and 6 % of interactions respectively. The three species differ strongly in their preferred habitat. *N. crassa* is a saprophytic fungus primarily living on burned vegetation in humid subtropical and tropical regions (Turner *et al.*, 2001).

*B. cinerea*, the grey mold-causing fungus, has a high host range including several vegetables like tomato, lettuce, and beans, or fruit crops such as strawberries and grapes (Chen *et al.*, 2023). Even though *B. cinerea* cause lesions on *Zea mays* under artificial conditions, cereals are not host plants of the pathogen (Bi *et al.*, 2023), thus it is unlikely that those fungi meet in the environment. In line, we never observed the outgrowth of new fungal hyphae from the contact point of intraspecies interactions, as it was reported for the interaction of *C. siamense* and *C. gloeosporioides* (Mehta and Baghela, 2021). Intriguingly, the two pathogenic species

*B. cinerea* and *C. graminicola* showed more stable fusion and germination frequencies as the saprophytic fungus *N. crassa*. This is in line with previous observations that especially plant-interacting species are able to tolerate wide ranges of ambient pH (Felix *et al.*, 1993).

In conclusion, our study has provided new insights into the molecular mechanisms underlying germling fusion in the maize pathogen *C. graminicola*. We have identified pH as a communication signal outgoing from secretomes harvested during the germling fusion, and have shown that the plasma membrane localized ATPase PmaI is a probable source for pH gradient formation. Our results also suggest that the a-pheromone receptor CgSte3 mediates sensing of acidic pH gradients, activating downstream the CWI MAPK pathway. These findings are consistent with previous studies that have shown that pH is a crucial environmental factor for fungi and its fluctuations influence diverse cellular processes. Our results also suggest that the maize pathogen *C. graminicola* communicates with the saprophyte *N. crassa* and the grey mold *B. cinerea*, although these three species differ strongly in their preferred habitat. Overall, our study contributes to a better understanding of the molecular mechanisms underlying germling fusion in fungi and indicate pH as fusion initiation signal across species borders.

## Acknowledgments

We thank Gabriele Beyer and Gertrud Stahlhut for excellent technical assistance and Lilith Elisabeth Gück and Simone Lewandowski for help with experimental work. This work was funded by the Deutsche Forschungsgemeinschaft (Bonn-Bad Godesberg). Grant was provided to D.E.N. (project NO 1230/3-1 (447175909). This work was partly supported by the Göttingen Graduate Center for Neurosciences, Biophysics, and Molecular Biosciences at the Georg-August-Universität Göttingen.

