## Supplementary Material for "PmaI-dependent pH gradients initiate inter- and intraspecies germling fusion in filamentous fungi"

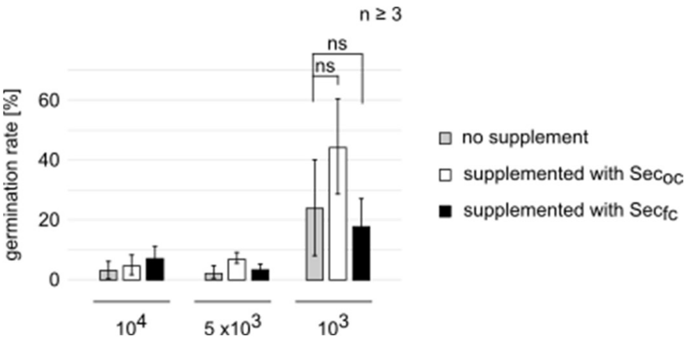


**Supplementary Figure 1** Germination rate of falcate conidia in dependence of spore concentration and the supplementation with 10 x concentrated $Sec_{oc}$ and $Sec_{fc}$. 10 µl of falcate conidia were dropped in different amounts ($10^4$, 5 x $10^3$, $10^3$) on water agar plates agar (1% serva agar, 1% agarose, 25 mM $NaNO_3$). After drying, the drops were topped with 20 µl of 10 x concentrated $Sec_{oc}$ and $Sec_{fc}$ and the liquid dried again. After 17 h incubation at 23°C, at least 100 conidia were microscopically examined for germination, n ≥ 3, ns = not significant.

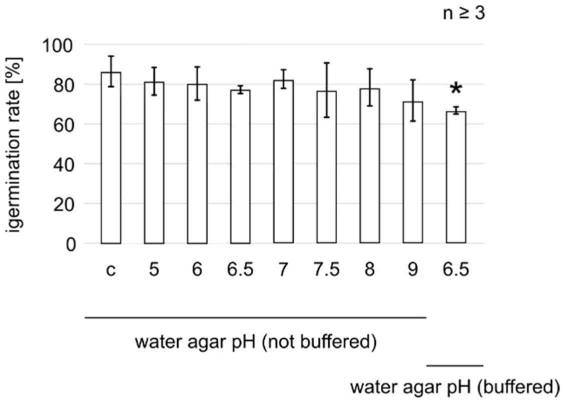


**Supplementary Figure 2** Germination rate of *C. graminicola* wild-type germlings derived from oval conidia in dependence on medium pH. Conidia were spread on water agar (1% serva agar, 1% agarose, 25 mM $NaNO_3$) adjusted to different medium pH with and without buffering. 50 µl of oval conidia (c = 5 x $10^7$ $ml^{-1}$) were spread on the corresponding medium plates. After 17 h of incubation at 23°C, at least 100 contacting germlings and conidia were microscopically examined for the formation of fusion bridges, n ≥ 3. Statistical significance was calculated with a two-tailed t-test corresponding to control conditions (c) *, p < 0.05.

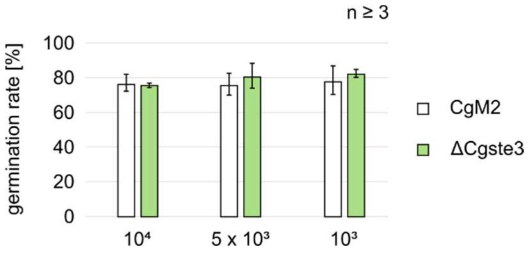


**Supplementary Figure 3** Spore concentration-dependent germination rates of *C. graminicola* Δ*Cgste3*. 10 µl of oval conidia spore solutions were dropped on water agar (1% serva agar, 1% agarose, 25 mM $NaNO_3$) to reach spore amounts of $10^4$ - $10^3$. Microscopic evaluation after 17 h of incubation at 23°C was performed for at least 100 conidia, n ≥ 3, statistical significance was calculated with a two-tailed t-test to CgM2 values of the corresponding spore concentration (*, p < 0.05).

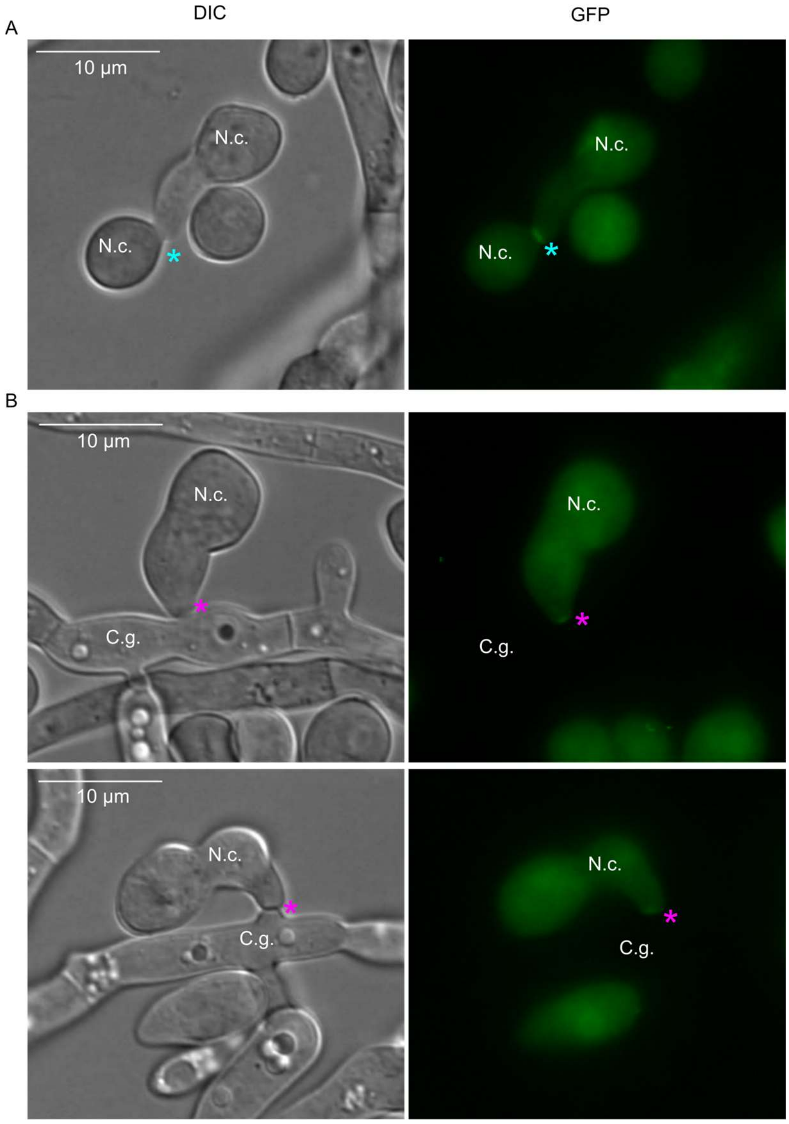


**Supplementary Figure 4** Mak-2-GFP localization in *N. crassa* at *C. graminicola* interaction sites. (A) 300 µl of *N. crassa* spore solutions (c = $10^7$ $ml^{-1}$) were spread on Vogels' MM and incubated for 3 h at 30°C. (B) 50 µl of a *C. graminicola* oval conidia spore solution (c = $5x10^7$ x $ml^{-1}$) was distributed on incubation medium and incubated for 14 h at 23°C. After that time, 300µl of Mak-2-GFP (Fleissner *et al.*, 2009) spore solution (c = $10^7$ $ml^{-1}$) was spread on those pre-incubated plates and further incubated at 28°C for 3.5 - 4 h. Mak-2-GFP localizations at *N. crassa* - *N. crassa* (A) and *N. crassa* – *C. graminicola* (B) interaction sites (light blue asterisks), scale = 10 µm.

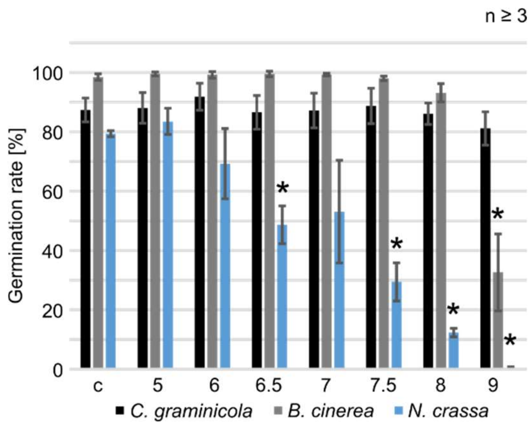


**Supplementary Figure 5** Germination rates of *C. graminicola*, *B. cinerea* and *N. crassa* on Vogel's minimal medium adjusted to different pH. Species-specific conidia amounts (*C. graminicola*: 50 µl of 5 x $10^7$ $ml^{-1}$; *B. cinerea* and *N. crassa*: 300 µl of c = $10^7$ $ml^{-1}$) were spread on Vogels' MM and incubated at 23°C for 17 h (*C. graminicola*, *B. cinerea*) or 28°C for 3,5 - 4 h (*N. crassa*), n ≥ 3, statistical significance was calculated with a two-tailed t-test to control conditions (c), *, $p < 0.05$.

**Supplementary Table 1** Fungal strains used in this study

| Organism | Strain | Genotype | Reference |
|---|---|---|---|
| *C. graminicola* | CgM2 (M1.001) | *C. graminicola* wild-type (wt) | (Forgey *et al.*, 1978) |
| *C. graminicola* | Δ*Cgso* | Homologous replacement of *Cgso* in CgM2, ssi, *hyg*$^R$, *Cgso*::*hph*, | (Nordzieke, 2022) |
| *C. graminicola* | Δ*Cgso::Cgso* | 5'*Cgso*::*Cgso*::3' *Cgso*, ssi, *nat*$^R$ | (Nordzieke, 2022) |
| *C. graminicola* | Δ*Cgste3* | Homologous replacement of *Cgste3* in CgM2, ssi, *hyg*$^R$, *Cgste3*::*hph*, | (Rudolph *et al.*, 2024) |
| *C. graminicola* | Δ*Cgste3::Cgste3* | 5'*Cgste3*::*Cgste3*::3' *Cgste3*, ssi, *nat*$^R$ | (Rudolph et al., 2024) |
| *N. crassa* | FGSC2489 | *N. crassa* mat A (wild-type) | FGSC |
| *N. crassa* | Wt::MAK-2-GFP | *his3*$^+$*::pef-1-mak-2-gfp, mat A* | (Serrano *et al.*, 2018) |
| *B. cinerea* | (B05.10) | *B. cinerea* wildt-type | (Quidde *et al.*, 1999) |

*nat*$^R$: resistant to nourseothricin; *hyg*$^R$: hygromycin resistant; ssi: single spore isolate, *hph*: hygromycin B phosphotransferase gene; FGSC: Fungal Genetic Stock Center
